# Numerous recursive sites contribute to accuracy of splicing of long introns in flies

**DOI:** 10.1101/290007

**Authors:** Athma A. Pai, Joseph Paggi, Karen Adelman, Christopher B. Burge

**Author notes:** These authors contributed equally (Athma A. Pai, Joseph Paggi).

## Abstract

Recursive splicing, a process by which a single intron is removed from pre-mRNA transcripts in multiple distinct segments, has been observed in a small subset of *Drosophila melanogaster* introns. However, detection of recursive splicing requires observation of splicing intermediates which are inherently unstable, making it difficult to study. Here we developed new computational approaches to identify recursively spliced introns and applied them, in combination with existing methods, to nascent RNA sequencing data from *Drosophila* S2 cells. These approaches identified hundreds of novel sites of recursive splicing, expanding the catalog of recursively spliced fly introns by 4-fold. Recursive sites occur in most very long (> 40 kb) fly introns, including many genes involved in morphogenesis and development, and tend to occur near the midpoints of introns. Suggesting a possible function for recursive splicing, we observe that fly introns with recursive sites are spliced more accurately than comparably sized non-recursive introns.

## Introduction

RNA splicing is a crucial step in the mRNA lifecycle, during which pre-mRNA transcripts are processed into mature transcripts by the excision of intronic sequences. Introns are normally excised as a single lariat unit after pairing of the 5’ and 3’ splice sites flanking the intron. However, some introns in the *Drosophila melanogaster* genome are known to undergo recursive splicing, in which two or more adjacent sections of an intron are excised in separate splicing events, each producing a distinct lariat [1,2]. Recursively spliced segments are bounded at one or both ends by recursive sites, which consist of juxtaposed 3’ and 5’ splice site motifs around a central AG/GT motif (with “/” indicating the splice junction) [1,3]. This mechanism appears to be restricted to very long *Drosophila* introns [3,4]. However, because recursive splicing yields an exon ligation product identical to that which would have been produced from excision of the intron in one step, the genome-wide prevalence and function of recursive splicing have been difficult to ascertain [3,4].

Recursive splicing was initially observed in the splicing of a 73 kb intron in the *Drosophila Ultrabithorax* (*Ubx*) gene, where the intron is removed in four steps through intermediate splicing of the 5’ splice site to two microexons and one recursive site before pairing with the proper 3’ splice site [1]. Bioinformatic searches for recursive sites predicted a couple hundred possible recursive sites in *Drosophila*, predominantly in introns larger than 10 kb [3], but sites in only four introns, all from developmentally important genes (*Ubx, kuzbanian* (*kuz*), *outspread* (*osp*), and *frizzled* (*fz*)), could be experimentally validated [1-3]. Biochemical characterization showed that recursive splicing is the predominant processing pathway for the splicing of these introns, which are generally constitutively spliced [1-4]. More recently, an analysis by Duff and coworkers of all ∼10 billion RNA-seq reads generated by the *Drosophila* ModENCODE project identified 130 recursively spliced introns in flies [4]. Using this larger catalog of recursive sites, they confirmed that recursive splicing is a conserved mechanism to excise constitutive introns, requires canonical splicing machinery, and only occurs in the longest 3% of *Drosophila* introns [4]. Similar analyses of mammalian RNA-seq datasets have resulted in the identification of just a handful of recursively spliced introns (4-5 introns), despite the greater abundance of long introns in vertebrate genomes, mostly in genes involved in brain development [5].

The scarcity of validated examples suggests that recursive splicing is quite rare, even in *Drosophila*. However, the transient nature of recursive splicing intermediates makes it difficult to detect evidence for recursive splicing using standard RNA-seq data. Support for recursive splicing has come from RNA-seq reads which span a junction between a known splice site and a putative recursive splice site in the middle of an intron, or from observation of a sawtooth pattern of reads resulting from the splicing out of a recursive segment [4,5]. Previous studies using polyA-selected RNA-seq data – which derive predominantly from mature transcripts – had limited ability to detect such evidence. However, nascent RNA sequencing, which profiles pre-mRNA transcripts shortly after they are transcribed, should enable much more efficient capture of reads from intermediates of splicing, including recursive splicing. Using such data should allow for more unbiased and systematic discovery of recursive splicing.

To globally detect transient splicing intermediates indicative of recursive splicing, we applied novel computational approaches to high-throughput sequencing data from short time period metabolic labeling. This approach detected about four times as much recursive splicing as had been previously observed. This expanded catalog of sites and associated analyses suggests a function for recursive splicing in improving splicing accuracy.

## Results

Pre-mRNA splicing can initiate immediately after transcription of an intron is completed, and can occur in as short a time as one or a few seconds [6-9]. Since recursive splicing involves the splicing of intermediate intronic segments, it may begin soon after the transcription of the first intronic recursive site. Thus, to have the greatest chance of capturing recursive intermediates, it is necessary to capture nascent transcripts shortly after transcription, before introns are fully spliced. Here, we used data from incorporation of a metabolic label to isolate RNA at short time points after transcription. The experimental approach to collect these data involved 5, 10, or 20 min labeling with 4-thio-uracil (4sU) in S2 cells, followed by RNA sequencing [9]. These data were complemented by steady state RNA-seq data representing predominantly mature mRNA (Methods).

We hypothesized that this high-coverage nascent RNA data would more readily identify recursive sites and better characterize the prevalence of recursive splicing. For this purpose, we used a computational pipeline to detect three key signatures of recursive splicing (Figure 1). First, we searched for splice junction reads derived from putative recursive sites (RatchetJunctions), as previously described (Figure 1A) [4,5]. Ratchet junction reads contain a segment adjacent to an annotated 5’ or 3’ splice site juxtaposed to a segment adjacent to an unannotated intronic recursive site, providing direct evidence for the presence of a recursive splicing event.

**Figure 1.**
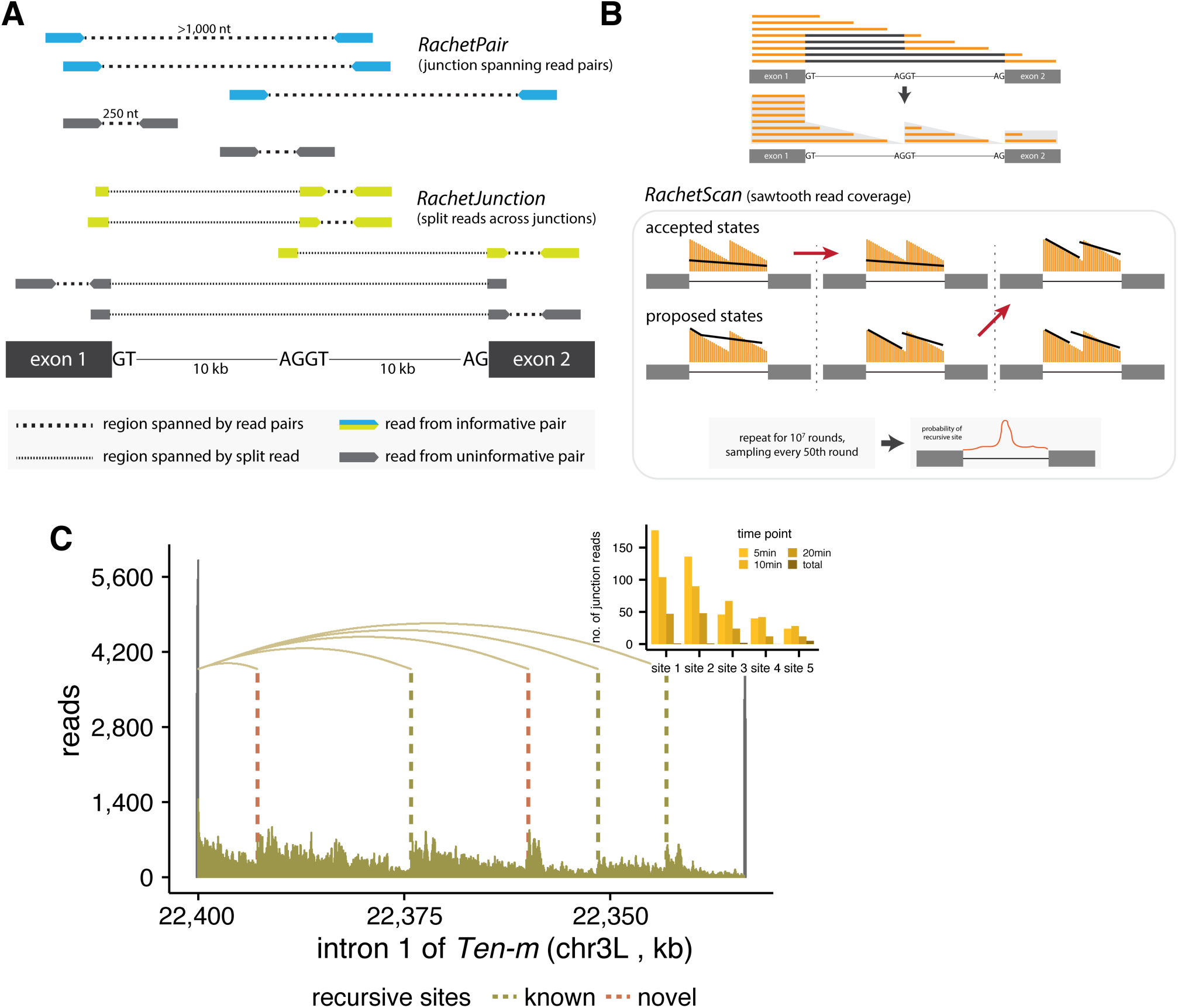
Identifying sites of recursive splicing using nascent 4sU-seq data. **(A)** Schematic indicating reads used for the RachetJunction (*bottom*) and RachetPair (*top*) methods to identify sites of recursive splicing. **(B)** Automated computational approach to detect sites of recursive splicing (*bottom*) using the sawtooth pattern created by co-transcriptional splicing of recursive segments (*top*). **(C)** Nascent RNA coverage across the first intron of *Ten-m*, which is recursively spliced. Vertical lines indicate location of detected recursive sites and curved lines indicate split junction reads between splice sites. The distribution of junction reads detected at each recursive site per timepoint is depicted in the inset.

Second, we developed a new computational tool, RatchetPair, to identify read pairs that map to distant genomic sites in a manner where presence of intervening recursive splicing can be inferred from the size distribution of inserts in the sequenced library (Methods; Figure 1A). Unlike ratchet junction reads, recursive junction spanning read pairs do not pinpoint a specific recursive site. Instead, a recursive site must be inferred based on the empirical distribution of fragment lengths and genomic sequence information. To do so, we adapted the GEM algorithm [10], originally designed to infer protein binding sites from ChIP-seq data, to assign a probability that each read-pair was indicative of a recursive site in a given region (Methods). This modified GEM algorithm was run with all read pairs and splice junction reads pooled together to derive the empirical distribution of fragment lengths.

Third, we developed the first automated software, RatchetScan, for inference of recursive sites from sawtooth patterns in read density (Figure 1B). This type of pattern is an expected product of co-transcriptional recursive splicing and has been associated with many recursive introns [4,5,11]. Briefly, assuming that RNA is spliced shortly after elongating past the transcription of the recursive splice site or 3’ splice site, the splicing of recursive segments during transcription of subsequent sequences will result in a sawtooth distribution of reads across the intron with recursive sites commonly located near the righthand base of each “tooth”.

RachetScan predicts the locations of recursive sites in three distinct steps. First, RNA-seq data was processed to summarize read density in each sub-intronic region (Figure 1 – figure supplement 1A). We then developed a Markov Chain Monte Carlo-(MCMC-) based inference algorithm to detect presence of sawtooth patterns in introns. This algorithm is suitable for efficient exploration of complex intronic read patterns encountered when considering a variable number of possible recursive splice sites in each intron. We were able to consider all nucleotides as potential recursive splite sites, rather than only focus on sites at the center of strong juxtaposed recursive motifs, allowing us to independently use sequence information to assess the false-positive rate of our method. Our *RachetScan* algorithm is initiated with a randomly chosen state, consisting of a set of proposed recursive sites in the intron (Figure 1 – figure supplement 1B). In each round, a new state is proposed by perturbing the current state, with three classes of perturbations: (1) a new recursive site is added; (2) a recursive site is removed; or (3) a recursive site location is locally shifted, each with defined probabilities. Using a scoring function and transition rules (detailed in Methods), the algorithm decides to either accept the new proposed state or maintain the current state. This procedure was iterated over 10^7^ rounds and the current state was sampled every 50 rounds, where the number of samples recorded in each state is proportional to the probability that the intron is best fit by the model corresponding to that state. Finally, recursive sites are predicted based on the output of the inference algorithm and sequence information (Figure 1 – figure supplement 1C; Figure 1 – figure supplement 2).

Combining these three approaches, our analysis detected 539 candidate recursive sites in 379 fly introns (Supplementary File 1). From this set, we curated a set of 243 “high confidence” recursive sites in 157 introns (with an FDR of 5%), and a “medium confidence” set of 296 sites (at an FDR of 20%; Figure 2A; Methods). Overall, 98 introns contained multiple recursive sites, with up to seven such high-confidence sites observed in a single intron. For instance, intron 1 of the tenascin major (*Ten-m*) gene contains five recursive sites, two of which were previously unknown (Figure 1C). Of the recursive sites previously reported by Duff and colleagues, 124 occurred in genes expressed in S2 cells. Our approach detected 119 (96%) of these known sites, as well as 126 novel high confidence sites and 294 novel medium confidence sites (Figure 2A), thus increasing the number of recursive sites defined in this cell type by ∼4-fold (Figure 1 – figure supplement 2C). Both the high confidence and the medium confidence candidate recursive sites exhibited a strong juxtaposed 3’/5’ splice site motif (Figure 2 – figure supplement 1). The greater numbers detected by our approach (2-4X more sites in this cell type), using less than 1/20^th^ as many reads as used by Duff and colleagues, confirms the potential of nascent RNA analysis for detection of recursive splicing.

**Figure 2.**
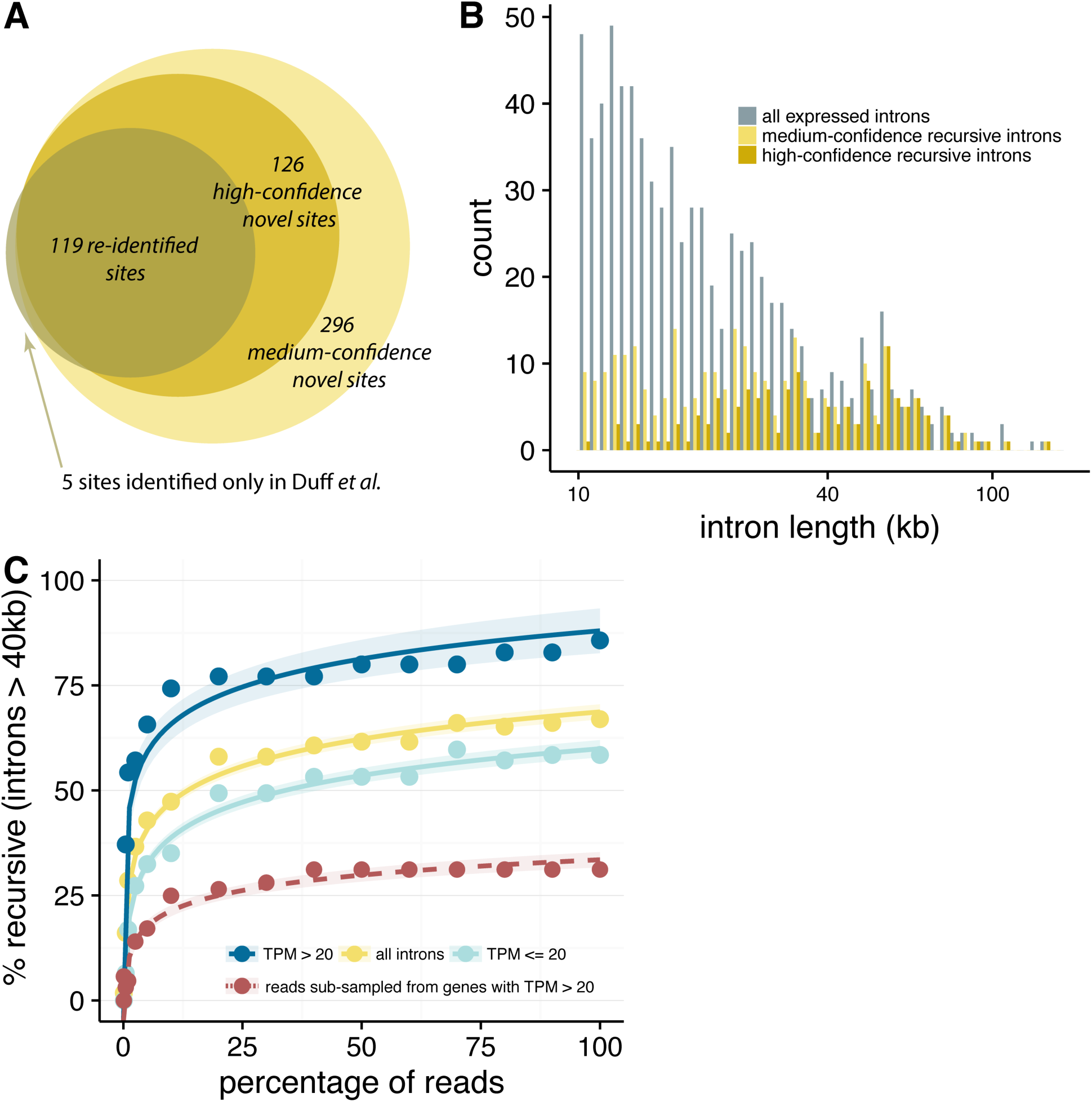
Recursive splicing is common mechanism to splice very long introns. **(A)** Number of recursive sites identified in this study, across sites previously identified, novel high-confidence sites, and novel candidate sites, with 5 sites that were previously identified but not detected in this study. **(B)** Distribution of intron lengths for all introns over 1kb (*grey*), with medium-confidence recursive introns in light tan and high-confidence recursive introns in dark tan. **(C)** Percentage of introns greater than 40 kb with at least one detected recursive site across various sub-samples of read coverage, where 100% indicates the percentage of recursive introns detected in the full dataset. Introns are subdivided into all introns detected (*yellow*), introns from lowly expressed genes (TPM ≤ 20; *light blue*), and introns from highly expressed genes (TPM > 20; *dark blue*).

Using this updated catalog of recursive sites, we observed that many very long introns (> 40 kb in length) have recursive sites, with 63% of such introns containing at least one high-confidence recursive site, and an additional 7% containing medium-confidence site(s) (Figure 2B). This observation suggests that recursive splicing is the prevalent mechanism by which very large fly introns are excised. We assessed the sensitivity of our detection pipeline by running it on subsamples of reads ranging from 0.1% to 100% of the total (Figure 2C). The shape of the resulting curve tapered off at higher coverage levels but never plateaued, indicating that new recursive sites were still being detected as read depth increased from 50% to 100% of sequenced reads and therefore would likely increase further at higher read depths. A somewhat higher proportion of recursive sites were detected in high-expressed genes (TPM > 20) than low-expressed genes (TPM ≤ 20). However, subsampling of the reads mapping to high-expressed genes to levels comparable to those observed for low-expressed genes resulted in a substantially lower fraction of recursive sites at each depth, suggesting that recursive splicing is more prevalent in low-expressed than high-expressed genes (Figure 2C). Together, these data suggest that the true fraction of very long introns that contain recursive sites may be substantially higher than our observed fraction of 63-70%, i.e. that recursive splicing is likely present in virtually all very long fly introns.

Recursive splice sites can be required for the processing of long introns [3]. However, it is possible that most recursive sites are functionally neutral, and that mRNA production is not impacted by their presence. The size of our dataset enabled us to examine four properties of recursive sites that could help to distinguish between these possibilities: sequence conservation; distribution in the fly genome; distribution within introns; and efficiency of splicing. In each case, the patterns observed suggest that recursive sites often have functional impact.

Both high and medium confidence recursive sites exhibited twice the level of evolutionary conservation observed in and around control AGGT motifs in long introns (Figure 3A), implying strong selection to maintain most or all of these sites. Recursively spliced introns were enriched in genes involved in functions related to development, morphogenesis, organismal, and cellular processes, with stronger enrichments for genes containing high-confidence recursive sites (Figure 3B; Supplementary File 2). Both of these observations are consistent with results from a previous study based on a smaller sample of recursive introns [4].

**Figure 3.**
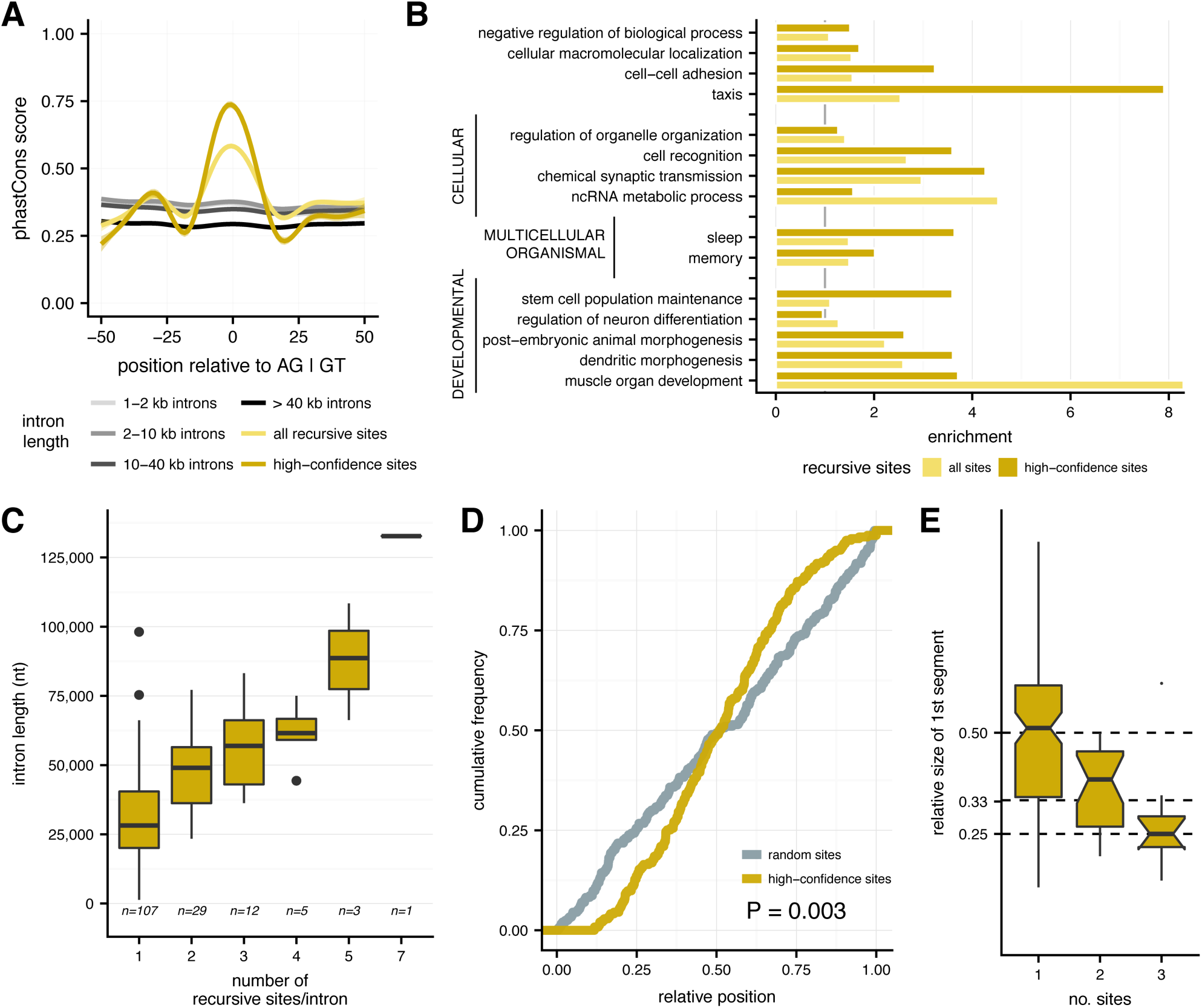
Characteristics of recursive sites in *Drosophila* introns. **(A)** Conservation of sequences around all detected recursive sites, with average phastCons scores for medium-confidence recursive sites (*yellow*), high-confidence sites (*gold*), and random AG|GT sites in introns increasingly larger than 1kb (*grey*). **(B)**. Enrichment of significant gene ontology categories among genes with any recursive site (yellow) and high confidence sites (gold), where gene ontology terms are broken down into their umbrella categories. **(C)**. Full intron length distributions for introns (*y-axis*) with varying numbers of recursive sites (*x-axis*). **(D)** Relative positions of recursive sites within introns for random sites chosen from a uniform distribution (*grey*) and single recursive sites in an intron (*dark tan*). **(E)** Distributions of the fractional distances (*y-axis*) of the first recursive segment for introns with increasing numbers of recursive sites (*x-axis*).

Longer introns might contain more recursive sites purely by chance. Indeed, while the majority of recursively spliced introns had just one recursive site, the number of sites increased roughly linearly with intron length (Figure 3C). However, the positioning of recursive sites within introns was significantly biased away from a random (uniform) distribution. Instead, recursive sites in introns with only one such site tended to be located closer to the midpoint of the intron than expected by chance (Kolmogorov-Smirnov *P* = 0.003; Figure 3D). Furthermore, the first recursive site in introns with two or three such sites tended to be located approximately 33% and 25% of the way from the 5’ end of the intron, respectively (Figure 3E). The distribution of recursive sites within introns suggests that they are positioned so as to break larger introns into “bite-sized” chunks of intermediate size (typically ∼9-15 kb in length) rather than at random locations which would more often produce much longer and much shorter segments. Recursively spliced introns were also enriched in first introns relative to subsequent introns in fly genes (hypergeometric P < 0.05).

To ask whether recursive splicing contributes to the efficiency of processing of very long introns, we evaluated the order and timing of recursive splicing events (Methods). We observed a steady increase in the proportion of exon-exon junction reads relative to recursive junctions across the time course, reflecting the progress of splicing (Figure 4A). Among recursive junction reads we observed far higher counts of reads spanning the 5’ splice site and the recursive site (RS), relative to RS-RS or RS-3’ splice site junctions, consistent with recursive segments being most often excised in 5’ to 3’ order (Figure 4A; Figure 4 – figure supplement 1B). This order of splicing is consistent with recursive splicing occurring cotranscriptionally.

**Figure 4.**
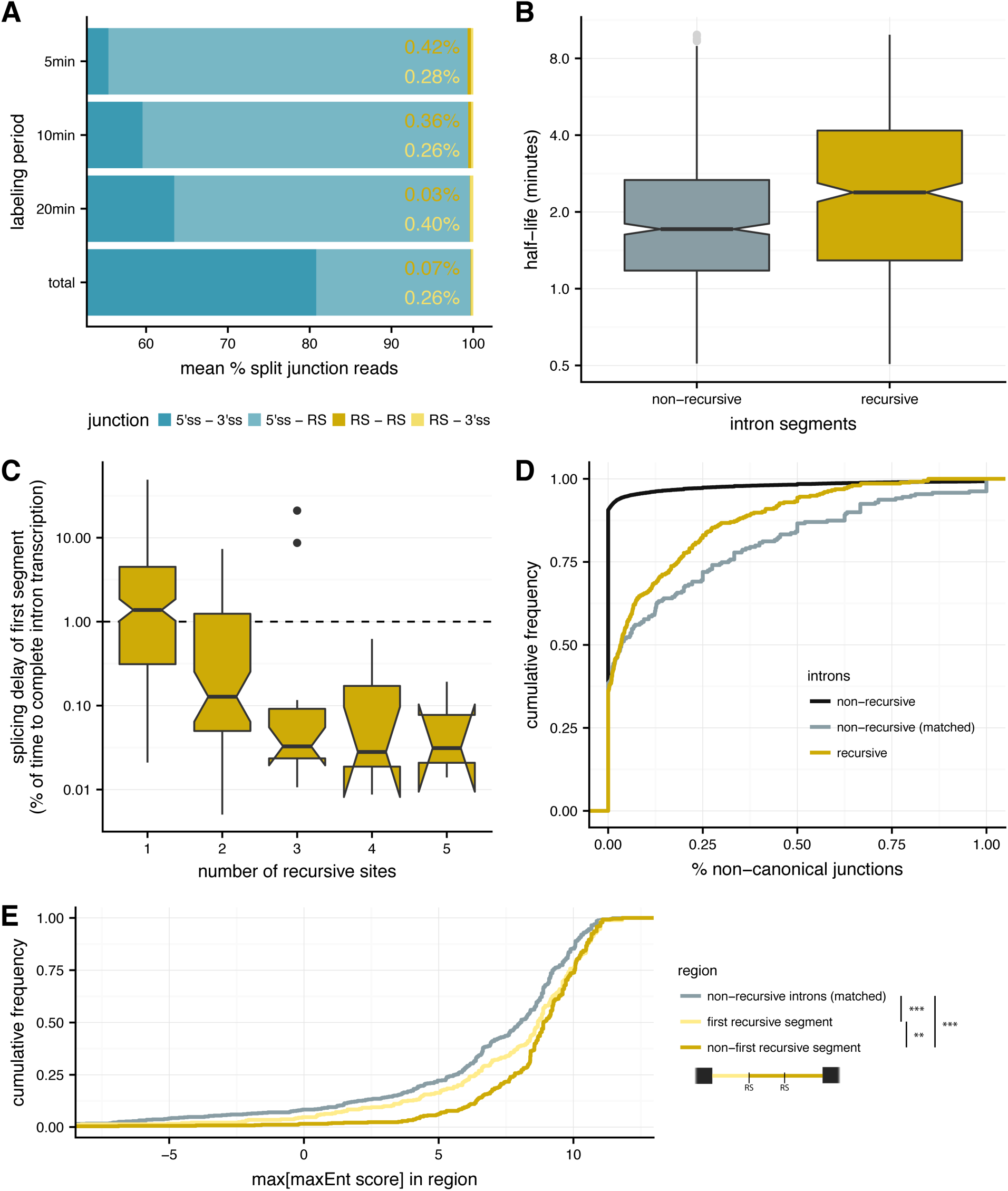
Recursive sites aid in efficient splicing of long *Drosophila* introns. **(A)** The percentage of splice junction reads (mean percentage across recursive introns; *x-axis*) that span the exon-exon boundary (5’ splice site – 3’ splice site; *dark blue*), a 5’ splice and recursive site (*light blue*), two recursive sites (*gold*), and a recursive site and 3’ splice site (*yellow*) across different labeling periods (*y-axis*). **(B)** Splicing half-lives for individual recursive segments (*dark tan*) and full non-recursive introns chosen to match recursive segment lengths (*grey*). **(C)** The delay in splicing half-life of the first recursive segment of an intron, relative to the time to transcribe the remainder of the intron (y-axis) for recursive introns with different numbers of recursive sites (x-axis). Increased splicing delays (> 1) are supporting of post-transcriptional splicing of the first segment, while decreased splicing delays (< 1) indicate co-transcriptional splicing. **(D)** Splicing accuracy measured by percentage of non-canonical unannotated reads for recursive introns (*gold*), non-recursive introns matched for intron length (*grey*), and all non-recursive introns (*light grey, dotted*). **(E)** Distribution of the highest maximum entropy score for an internal 5’ splice site motif (*x-axis*) for initial recursive segments (not expected to contain an RS-exon; *yellow*); non-first recursive segments (potentially contain RS-exons; *gold*), and non-recursive introns matched for intron length (*grey*). Significance is indicated such that **: P < 0.01 and ***: P < 0.001, with a Kolmogorov-Smirnov test.

Previously we developed a framework for estimating rates of splicing from nascent RNA sequencing data across different labeling periods [9]. Here, we adapted this approach to estimate the splicing half-lives of individual recursive segments (Methods; Figure 4 – figure supplement 1A; Supplementary File 3), which have a mean length of 9.1 kb (Figure 4 – figure supplement 1C). Recursive segment half-lives were the slowest for the first segment in the intron, with faster half-lives for successive segments (Figure 4 – figure supplement 1D). Overall, recursive segments had 1.5-fold longer half-lives than non-recursive introns of the same lengths (Figure 4B; Mann-Whitney *P* = 1.5 × 10^−9^). Estimating the mean splicing half-life of a recursive intron as the maximum of a set of exponentials (to approximate the waiting time to splice all recursive segments), we found that recursive introns are spliced more slowly than non-recursive introns of similar size (Figure 4 – figure supplement 1E; Mann-Whitney *P* < 2.2 × 10^−16^), consistent with the larger number of biochemical steps involved in recursive versus non-recursive splicing of an intron.

To ask whether recursive splicing occurs while the intron is continuing to be transcribed, we calculated the ratio of the half-life of the first segment to the estimated time needed to transcribe the remainder of the intron (Methods). For 49% of recursively spliced introns, the first segment half-life is shorter than the time to transcribe the full recursive intron (Figure 4C), implying common co-transcriptional splicing in about half of cases. We observed that longer recursive introns were more likely to be spliced co-transcriptionally.

The accuracy of splicing is likely to be at least as important as its speed, since splicing to an arbitrary (incorrect) splice site will most often produce an mRNA that is unstable or encodes a protein that is aberrant or nonfunctional. As a simple measure of potential splicing errors, we tallied the fraction of reads that spanned “non-canonical” splice junctions, involving pairs of intron terminal dinucleotides other than the three canonical pairs “GT-AG”, “GC-AG” and “AT-AC” that account for ∼99.9% of all known fly introns. For the bulk of non-recursive introns (most of which are < 100 nt in length), the frequency of such non-canonical splicing was negligible (Figure 4D, black curve). However, for non-recursive introns with lengths matching the much more extended lengths of recursively spliced introns, potential splicing errors were much more frequent (Figure 4D, gray curve), suggesting that the fly spliceosome loses accuracy as intron length (and the number of possible decoy splice sites) increases. Notably, recursive introns had ∼37% fewer non-canonical junctions compared to similarly sized non-recursive introns (Figure 4D, gold curve, Kolmogorov-Smirnov *P* = 0.015). Therefore, presence of recursive splice sites may increase the accuracy of splicing, perhaps at the expense of splicing speed.

## Discussion

Analysis of intermediates can provide insight into otherwise hidden biochemical pathways. Here, application of new computational approaches to nascent RNA sequencing data, which is highly enriched for splicing intermediates, enabled us to identify about four times more recursive sites in the *Drosophila* genome than were known previously. The surprisingly widespread occurrence of recursive splicing raises questions about what functions it may serve.

A priori, this pathway might improve the speed or accuracy of splicing, or might impact regulation. Our analyses suggest that recursive splicing does not in fact increase splicing rates, and may actually slow splicing somewhat, likely because of the additional steps involved.

However, we observe that the *Drosophila* splicing machinery appears to make a relatively high rate of errors in the splicing of longer introns, and that presence of recursive sites may substantially improve splicing accuracy. In splicing of a non-recursive 30 kbp intron, the 5’ splice site is synthesized about 20 minutes before it’s correct partner 3’ splice site, creating a long window during which splicing can only occur to incorrect 3’ splice sites, likely contributing to the higher error rate seen for long fly introns. Presence of a recursive site may help to organize the processing of the intron, keeping the splicing machinery associated with the 5’ splice site engaged in a productive direction and avoiding involvement with decoy 3’ splice sites. It was previously observed that masking a recursive splice site in a zebrafish *cadm2* intron does not change the overall splicing of the intron but reduces *cadm2* mRNA levels [5]. This observation could be explained if the recursive site promotes accurate splicing and prevents unproductive splicing pathways that result in unstable products targeted by RNA decay pathways such as nonsense-mediated mRNA decay.

Recursive sites may also participate in splicing regulation. A previous study of a handful of recursively spliced introns in humans identified RS-exons that are initially recognized during recursive splicing following an “exon definition” model of splice site recognition [5], while an alternative “intron definition” pathway has been proposed for recursive splice site recognition in flies [4]. An exon definition model would require presence of a 5’ splice site downstream of each recursive site. Consistent with this model, we observed that recursive segments following recursive sites are enriched for strong 5’ splice site motifs relative to first recursive segments and to non-recursive introns matched for length (Figure 4E). Use of an exon definition pathway in the initial steps of spliceosome assembly might also contribute to splicing accuracy, with the downstream 5’ splice site helping to specify the recursive site [9]. It could also produce alternative mRNA isoforms containing an additional exon [5].

Exon definition of recursive segments through transient RS-exons requires that the recursive site first be recognized as a 3’ splice site and subsequently as a 5’ splice site for splicing of the subsequent segment (assuming that simultaneous recognition of an RS in both modes is sterically prohibited). For this ordered recognition to occur (and for sequential splicing of recursive sites generally), binding of dU2AF/U2 snRNP must outcompete binding of U1 to the RS prior to its splicing to the upstream exon. Consistent with this expectation, the 3’ splice site motifs of RS are very strong, stronger than non-recursive 3’ splice sites, and they have higher information content than RS 5’ splice sites (Figure 4 – figure supplement 2).

Why are developmental genes enriched for long introns with recursive splicing? No clear answer emerges. It is possible that intron length is used to tune the timing of expression of these genes relative to the rapid embryonic cell cycle [12,13]. Alternatively, long introns may be needed to accommodate large transcriptional enhancers or complex three-dimensional organization of these gene loci related to their dynamic transcriptional regulation, or to facilitate alternative splicing. In addition to producing unstable mRNAs, splicing errors may also produce stable mRNAs that encode aberrant protein forms, including dominant negative forms. Perhaps recursive splicing has been selected for in these genes to improve splicing accuracy and avoid production of aberrant developmental regulatory proteins at critical stages to improve the robustness of development.

## Methods

### RNA-seq data analysis

We used RNA-seq data from our recent study of splicing kinetics in *Drosophila* S2 cells (GEO GSE93763; [9]). These data included 3 independent replicates of S2 cells labeled for 5, 10 and 20 minutes with 500 μM 4-thiouridine, isolation of labeled RNA, and library preparation using random hexamer priming following ribosomal RNA subtraction. cDNA for two independent biological replicates of ‘total’ RNA were prepared using an equal mix of random hexamers and oligo-dT primers from unlabeled S2 cells [9]. Libraries were sequenced with paired-end 51 nt reads (100 nt reads for the ‘total’ RNA samples), generating an average of 126M read pairs per library. Reads were filtered and mapped to the *Drosophila melanogaster* dm3 reference assembly as described in [9].

Gene expression values (TPMs) in each replicate library were calculated using Kallisto [14] and the transcriptome annotations from FlyBase *Drosophila melanogaster* Release 5.57 [15].

### Identifying sites of recursive splicing

We used three features of recursive sites found in our nascent sequencing data to identify recursive sites: (1) splice junction reads derived from putative recursive sites (“RachetJunctions”), (2) recursive-site spanning pairs, specifically read pairs that map to sites flanking putative recursive segments such that the fragment length can only be accounted for by the presence recursive intermediate (“Rachet Pair”), and (3) a sawtooth pattern in intronic read density (“RachetScan”). Details of the computational and statistical methods for each of these approaches and our pipeline for recursive site detection are described below.

Out of the full set of recursive sites that were identified across all three methods, we filtered down to a final set of sites with the following criteria: (1) in genes with TPM ≥ 1 in the total RNA libraries, (2) in introns with at least 3 reads spanning the 5’ to 3’ splice sites (using the largest annotated intron), and (3) not overlapping with an annotated 5’ splice site in the that intron. This resulted in a total of 539 recursive sites identified by any method. High-confidence sites were identified by the criteria used by Duff *et al*. [4]. We wrote a script to plot the read density around putative recursive sites and manually filtered each site based on the presence of a recognizable sawtooth pattern. This resulted in the identification of 243 high-confidence sites.

Conservation of recursive sites was estimated using per nucleotide phastCons scores [16] from a 15-way *Drosophila* alignment downloaded from UCSC Genome Browser.

#### RachetJunction identifying splice junction reads from recursive intermediates

Splice junction reads that span putative recursive junctions provide direct evidence for recursive splicing (Figure 1A bottom). In order to identify such reads, we extracted the coordinates of annotated introns and exon-exon junctions from FlyBase D. melanogaster Release 5.57 and aligned the 4sU-RNAseq reads to the corresponding genome release using hisat2 [17]. We then used pysam [18] to extract reads with an upstream junction matching an annotated 5’ splice site and a downstream end mapping to an AGGT that is upstream of the downstream most corresponding annotated 3’ splice site.

#### RachetPair identifying recursive-site spanning pairs

In addition to splice junction reads, read pairs with one end on either side of a recursive splice junction – henceforth referred to as recursive junction spanning read pairs – provide evidence for recursive sites. We defined putative recursive junction spanning read pairs as read pairs with a first read aligning close upstream of an annotated 5’ splice site and a second read aligning to an intronic region more than 1000 nt downstream of the first read. Additionally, we filtered out read pairs than have an insert length of less than 1000 nt conditioned on completion of an annotated splicing event (excluding cassette exons with an AGGT at their 5’ end).

Unlike splice junction reads, recursive junction spanning read pairs do not immediately implicate a specific recursive site. Instead, a recursive site must be inferred based on the empirical insert length distribution and genomic sequence information. To do this, we adapted the GEM algorithm, which was originally used to infer protein binding sites from ChIP-seq data [10].

Our modifications to the algorithm and choices for parameters described in Guo *et al*. are as follows:

1. The probability of a read, *r*_*n*_, given that there is a recursive site at position *m, P(r*_*n*_|*m)*, was defined as the probability of observing the implied insert length in the empirical insert length distribution.
2. The prior probabilities of each position being a recursive site, Π_1 − *N*_, were set such that Π_*j*_ ∝ *max* (0,*M*(*i*) − 0.8), where *M(i)* is the motif score for position *i* as described above. This function was used to determine the prior probabilities that reflect the preference for strong motifs observed in the Duff *et al*. set of recursive sites [4].
3. Recursive splice junction reads were counted within the number of effectively assigned reads in the M-step. This ensured that sites with support from recursive junction reads are more likely to be recursive sites.
4. The sparsity parameters, *α*_4_, was defined as the number of assigned reads divided by 40.
5. The algorithm converged when prior probability did not change by more than 10^−5^ between iterations. Upon convergence, read pairs were assigned to a putative recursive site using the MAP estimate.

The modified GEM algorithm was run with all read pairs and splice junction reads pooled together.

#### RachetScan identifying sawtooth pattern in recursive intron read density

Recursively spliced introns contain a distinct “sawtooth” pattern due to the co-transcriptional nature of splicing. This is depicted in Figure 1B, where the horizontal lines represent elongating pre-mRNAs – with a uniform distribution of elongation distances across a population of cells over time – and the blacked out sections represent segments that have already been spliced out and degraded. Reads sequenced from the nascent RNA population will only be derived from the sections of RNA that have not yet been spliced and degraded, such that their density across the intron exhibits linear decay across each recursive segment.

We developed an algorithm to predict recursive splice sites from the presence of a sawtooth pattern in introns. Our algorithm consists of three distinct phases: pre-processing of the RNA-seq data, Monte Carlo Markov Chain based inference of the presence of a sawtooth pattern, and the prediction of recursive sites based on the output of our inference and sequence information.

#### RNA-seq read pre-processing (visualized in Figure 1 – figure supplement 1A)

We searched for the presence of a sawtooth pattern in the read distribution of all introns over 8 kb that had at least one spanning splice junction read in any sample. Empirical testing suggested our method displayed a high rate of false positives in introns under 8kb, likely due to regression over short segments being more sensitive to noise in read density. We removed regions annotated as exons using bedtools `subtract`. The number of read pairs aligning to each position were summed to obtain per base coverage counts, where read pairs straddling a given position were counted as a positive alignment.

In order to avoid erratic read coverage in repeat regions inhibiting our ability to perform meaningful regressions in later steps of the analysis, we masked the read densities in repeat regions and replaced the read counts in RepeatMasker annotate repeat regions [19] and the 100 flanking nucleotides with the median read density from the 900 nt flanking either side. This length was chosen because it was short enough that read densities in this range were comparable to those in the masked region, but long enough to avoid sensitivity to noise in read densities. To attain additional smoothing and reduce the time required to perform the regressions in the next step of our analysis, we separated introns into 100 nt bins and calculated the average of each bin. Throughout the rest of our analysis, we represented the read density of each intron using arrays of these average values.

#### Regression

We performed linear regression on all sub-regions of each intron. We assumed that variance in read density at each position was proportional to the coverage level at that position, which is likely true since RNA-seq read coverage is intrinsically the sum of Bernoulli random variables. To calculate these regressions, we developed a function that made use of the Scipy stats weighted linear regression function [20] as a sub-process, such that:

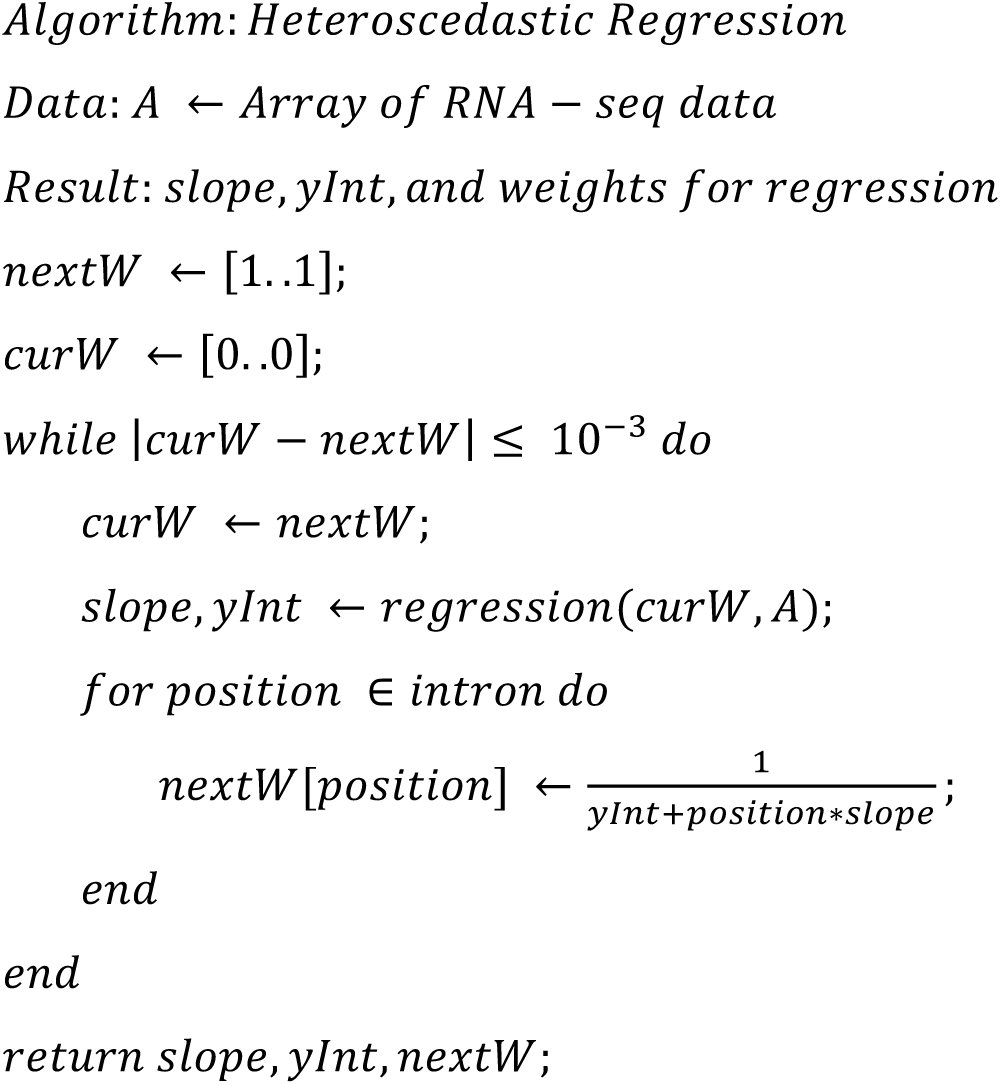

Note that | *curW* − *nextW* | ≤ 10^−3^ checks whether all weights have changed by at most 10^−3^.

#### Monte Carlo Markov Chain (MCMC; visualized in Figure 1 – figure supplement 1B)

We developed a Monte Carlo Markov Chain (MCMC) algorithm to detect the presence of a sawtooth pattern in each intron. Our algorithm is round based such that upon entering each round, we have an accepted state consisting of a set of proposed recursive sites in the intron. In each round, a new state is proposed by perturbing the current state. We use a scoring function and transition rules (defined below) to decide if we wish to accept this proposed state or continue with the current state. This procedure is iterated for 10^7^ rounds and a sample of the current state is recorded every 50 states. The number of samples recorded in each state is proportional to the probability that the intron is best fit by the model corresponding to that state. Therefore, to attain probabilities that each state is the most accurate model, we normalize the number of samples recorded in each state by the total number of samples.

There are three classes of perturbations used to propose new states (depicted in Figure 1 – figure supplement 1B):

1. A new recursive site was added probabilities 0.4 (visualized as transition 1 & 2).
2. A recursive site was removed with probability 0.4 (visualized as transition 4).
3. A recursive site was slightly perturbed with probability 0.2 (visualized as transition 3).

States are scored using a function taking into account how well the corresponding regression fits the observed RNA-seq read density as well as the number of free parameters in the model. The scoring function is based on the Bayesian Information Criteria (BIC), such that:

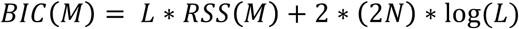

where *RSS*(*M*) is the weighted sum of squared deviations for all recursive segments, *L* is the intron length, and *N* is the number of recursive sites. Note that 2*N* is the number of free parameters in the model, as each recursive segment is fit for its own slope and y-intercept. The score is then given by:

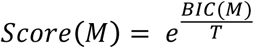

where T is a constant used to scale the magnitude of the scores. T = 5 was used for all analyses presented here. In order to constrain our algorithm to fit sawtooth patterns and note more general patterns in the read density, new states are only considered if, at each recursive site, the RNA-seq density predicted by regressions increased by at least 1.5 fold.

We use the standard transition rules for MCMC inference, which we outline here for convenience. If the score for the new state is lower than the score for the old state, the new state is deterministically adopted. Otherwise, the new state is adopted with probability score_new_/score_old_. When the old state had zero recursive sites, this probability was divided by 2 to account for the imbalance in transition probabilities. We chose parameters for burn-in-time, number of iterations and sampling frequency that empirically resulted in consistent convergence across multiple runs of the algorithm. These values were: a burn in of 10^5^ iterations, sampling frequency of 50 iterations, a total of 10^7^ iterations.

After all samples were collected, we calculated the probability that each position in the intron is a recursive site. For each position, we summed the occurrences of that position as a recursive site across all samples. Probability scores were then calculated for each position by dividing this sum by the total number of samples.

#### Peak Calling (visualized in Figure 1 – figure supplement 1C)

We predicted recursive sites from the MCMC probability scores in a two step process. First, regions with probability above a given threshold (0.08) were recorded. Any of these regions within 500 nt of each other were merged. For each of these regions, a position potential function, P, was defined as 1 inside the peak and flanked by a logistically decaying curve on either side. The logistic function is given by:

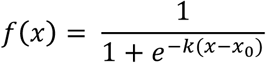

The parameters were set as *x* _0_= 500 nt from either end and k was set as 6/500 for the left flank and −6/500 for the right flank. The resulting distribution has values very close to zero at 1000 nt away from the peak and values of 0.5 at a distance of 500 nt. This distribution was chosen based on the empirical performance of the MCMC-based inference when compared to random. Each AGGT in the intron was then scored by the following equation and the maximum scoring AGGT was then reported as a putative recursive site:

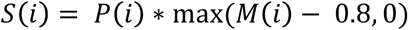

#### FDR Quantification

Shuffled peaks were produced to evaluate the false discovery rate of the sawtooth pattern identification pipeline. For each intron, the initially recorded regions of probability exceeding 0.08 were redistributed with uniform probability across the intron. The length and number of regions were maintained. The remainder of the peak calling procedure was then applied to obtain an null distribution of recursive probability peaks.

### Motif Scoring

We calculated position weight matrices (PWM) for the intronic portions of Drosophila 5’ and 3’ splice sites using all annotated splice sites. These weight matrices were then juxtaposed with the 3’ splice site PWM followed by the 5’ splice site PWM to create a recursive splice site motif PWM. Individual motif occurrences were scored using a normalized bit score [21]. The bit score for each motif occurrence is defined as the sum across the log probabilities for each nt being drawn from the motif. We calculated normalized scores by subtracting the minimum possible score and dividing by the range of possible bit scores.

### Estimating the true number of recursive sites

In order to assess the sensitivity of our recursive site detection pipeline, we subsampled our reads to various proportions of the total read coverage and re-assessed the number of recursive sites detected. To do so, we used the *samtools view* − *s* command [22] to subsample each fastq file from all samples to the following fractions: 0.1%, 0.5%, 1%, 2.5%, 5%, 10%, 20%, 30%, 40%, 50%, 60%, 70%, 80%, and 90%. For each of these subsampled read sets, we re-ran the entire recursive site detection pipeline as described above to assess the number of recursive sites detected.

To assess the impact of gene expression levels on our power to detect recursive sites, we separated long introns into those from lowly expressed genes (TPM ≤ 20) and highly expressed genes (TPM > 20). Using the subset of reads mapping to these genes, we repeated the subsampling procedure and entire recursive site detection pipeline described above to characterize the percentage of lowly or highly expressed long introns that have recursive sites.

Finally, to understand whether the lower proportion of lowly expressed introns that have recursive sites is due to technical or biological reasons, we subsampled reads from long introns within highly expressed genes to match the read distribution of a comparable number of long introns from lowly expressed genes. Specifically, we isolated all reads from long introns in highly expressed genes and used pysam [18] to randomly subsample these reads to match the distribution of reads from lowly expressed introns. Using only this subset of reads from highly expressed genes, we again repeated the subsampling procedure and the entire recursive site detection pipeline described above to characterize the percentage of highly expressed introns that have recursive when reads from these introns are subsampled to a lower read coverage.

### Determining the order of recursive splicing

Previous studies have searched exclusively for recursive junction reads consistent with the 5’ to 3’ removal of recursive segments [4,5]. In order to determine if recursive splicing does indeed follow a 5’ to 3’ order, we quantified junction reads consistent with alternative orders of recursive splicing. These reads fall into two categories: junction reads between two intronic AGGTs and junction reads from an intronic AGGT to an annotated 3’ splice site.

We constrained our search to combinations or recursive sites producing recursive segments of at least 1 kb. Nearly all recursive segments detected in our study were greater than 1 kb, thus adding this constraint mainly served to filter out spurious hits likely caused by alignment errors and unannotated splicing events. We considered all events with support from at least 3 uniquely aligning reads with recursive splice sites scoring above 0.85 in the scoring metric described above. Requiring at least three uniquely aligning reads matches the cutoff used for our previous analysis, where we found that recursive splice sites generally have strong motifs that score greater than 0.85.

These analyses produced thirteen candidate intronic AGGT to annotated 3’ splice site recursive junction reads, and no candidate intronic AGGT to AGGT recursive sites. These candidate recursive splice sites were evaluated visually in a genome browser. Two of these sites corresponded to recursive splice sites detected by both methods in our study. One of these sites has sixty recursive junction reads supporting a 5’ to 3’ order, while only five junction reads supporting a 3’ to 5’ order. The second site has 829 and 13 junction reads for the 5’ to 3’ and 3’ to 5’ orders, respectively. All other candidate alternative ordering sites did not appear to be true recursive sites, due to either a lack of sawtooth pattern, low intron expression, or extensive repeats complicating the alignment. These data suggest that recursive splicing overwhelmingly, but perhaps not always, proceeds in a 5’ to 3’ order.

### Splicing rates in recursively spliced introns

We quantified splicing rates for each recursive segment independently by applying an approach for 4sU RNA-seq data that we previously described [9]. Specifically, we used reads that overlapped recursive sites and junction reads (split between either the recursive site and an annotated splice site, between two recursive sites, or between the 5’ and 3’ splice sites; as detailed in Figure 4 – figure supplement 1A), as measures of uncompleted and completed segment splicing, respectively. The junction dynamics approach from Pai *et al*. 2017 [9] was applied to each set of reads to obtained a splicing half-life for each recursive segment. For full introns matched for length, we used splicing half-lives calculated in Pai *et al*. 2017 [9]. We estimated co-vs. post-transcriptional splicing of the first recursive segment by comparing the segment splicing half-life to the time to transcribe the remainder of the intron. Specifically, the time to complete intron transcription was estimated as:

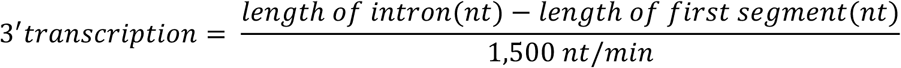

and the splicing delay was calculated as the ratio of the first segment’s splicing half-life to the 3’ transcription time.

#### Estimating the rate of splicing of full recursive introns

To estimate the mean lifetime of a recursively spliced intron, we estimated the waiting time for all recursive segments to be spliced out by calculating the maximum of the set of individual exponentials from each segment. For one exponential, the mean lifetime is 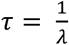 where *λ* is the coefficient from the exponential fit. There is an analytical solution for estimating the mean lifetime in situations where there are only two exponentials to be combined. Thus, we limited our analysis to recursive introns with only one recursive site, corresponding to the presence of two recursive segments (*i*.*e*. two exponentials). For these introns, the mean lifetime *τ*_*recursive*_ can be calculated by:

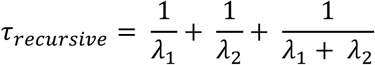

where *λ*_*1*_ is the exponential coefficient for the first segment and *λ*_*2*_ is the exponential coefficient for the second segment. To conservatively compare our recursive intron *τ*_*recursive*_ values with the mean lifetimes of non-recursive introns, we added the time necessary for the first segment to be transcribed to our *τ*_*recursive*_, with the rationale that the first segment must be completely transcribed before the second can begin to be spliced. Assuming a 1.5 kb/min transcription rate 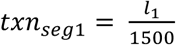, where *l*_1_ is the length of the first segment (in nucleotides).

### Estimating splicing accuracy

We estimated the accuracy of splicing in *Drosophila* introns by identifying non-annotated junction reads with non-canonical splice site sequences within annotated introns. To do so, we first re-mapped the raw 4sU-seq reads with the STAR v2.5 software [23], with the mapping parameter `--outSAMattribute NH HI AS nM jM` to mark the intron motif category for each junction read in the final mapped file.

The `jM` attribute adds a `jM:B:c` SAM attribute to split reads arising from exon-exon junctions. All junction reads were first isolated and separated based on the value assigned to the `jM:B:c` tag. Junction reads spanning splice sites in the following categories were considered to be annotated or canonical: (1) any annotated splice site based on FlyBase *D*. *melanogaster* Release 5.57 gene structures [`jM:B:c,[20-26]`], (2) intron motifs containing “GT-AG” (or the reverse complement) [`jM:B:c,1` or `jM:B:c,2`], (3) intron motifs containing “GC-AG” (or the reverse complement) [`jM:B:c,3` or `jM:B:c,4`], and (4) intron motifs containing “AT-AC” (or the reverse complement) [`jM:B:c,5` or `jM:B:c,6`]. Junction reads with `jM:B:c,0` were considered to arise from non-canonical non-annotated splice sites. We calculated the frequency of inaccurate splice junctions for each intron as a ratio of the density of reads arising from non-canonical non-annotated splice sites to the density of all junction reads from the intron.

### Calculating splice site scores

We calculated the strength of splice sites using a maximum entropy model as implemented in maxEntScan [24] using 9 nucleotides around the 5’ splice site (−3:+6) and 23 nucleotides around the 3’ splice site (−20:+3). These models were optimized on mammalian splice site preferences, but seem to be reasonable for *Drosophila* as well and have been used in gene prediction in fly genomes.

### Gene Ontology analyses

Gene Ontology enrichment analyses were performed using a custom script to avoid significant gene ontology terms with overlapping gene sets. Specifically, the script used the Flybase gene ontology annotation downloaded from the Gene Ontology Consortium website [25] and searches for the gene ontology term with the most significant enrichment of genes with recursively spliced introns (relative to a background of all genes with introns greater than 10,000 kb). Genes that belong to the most significant gene ontology term are then removed from the foreground and background sets of genes and the process is repeated iteratively until no genes are left in the foreground set. P-values are computed using a Fisher-exact test and then corrected using a Benjamini-Hochberg multiple test correction.

### Code availability

Source code is available upon request and will be made available in a public repository prior to publication.

## Acknowledgements

We thank members of the Burge lab, Brent Graveley and Phil Sharp for helpful comments on this manuscript.

## Author contributions

Athma A Pai, Software, Formal analysis, Investigation, Visualization, Methodology, Writing – original draft;

Joseph Paggi, Software, Formal analysis, Investigation, Visualization, Methodology, Writing – review and editing;

Karen Adelman, Supervision, Funding acquisition, Writing – review and editing;

Christopher B Burge, Conceptualization, Supervision, Funding acquisition, Writing – original draft

## Supplementary Figure Legends

**Figure 1 – figure supplement 1.**
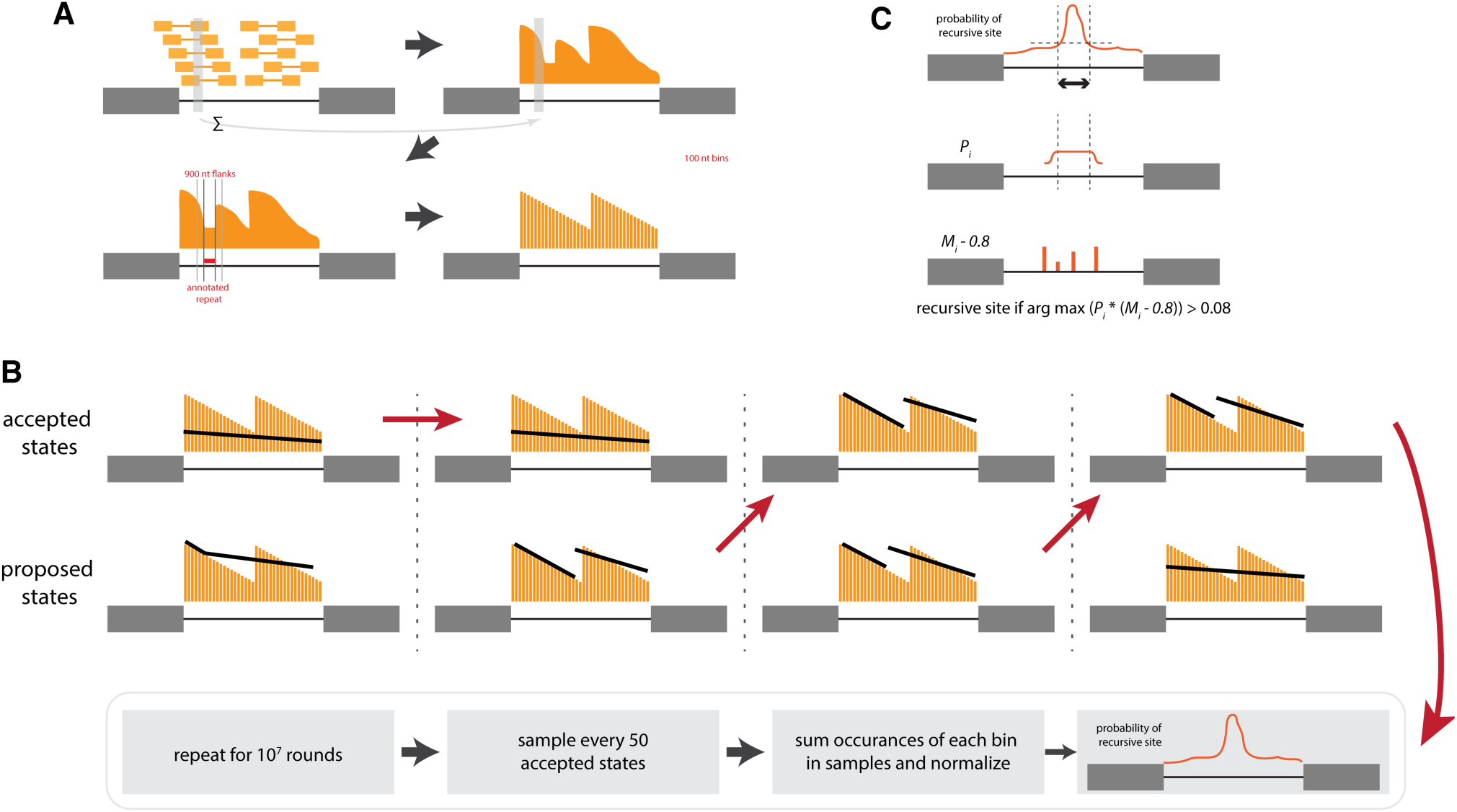
RachetScan method for automated detection of sawtooth patterns indicating of recursive splicing. **(A)** RNA-seq pre-processing steps to convert reads into an array of read densities: (1) summing the read coverage for each base-pair (*top*) and (2) replacing the read counts in annotated repeat regions and 100 flanking nt with median read density in 900 nt flanking regions (*bottom*) **(B)** MCMC algorithm infers probability that each position in intron is a recursive splice site, where upon entering each round with a previously accepted state, this state is perturbed to propose a new state and the new state is either accepted or rejected. The procedure is performed over 10^7^ rounds, with sampling every 50 rounds to obtain a probability that each base pair is a recursive site. **(C)** Sequence information is used in conjunction with MCMC-inferred probabilities to predict recursive sites.

**Figure 1 – figure supplement 2.**
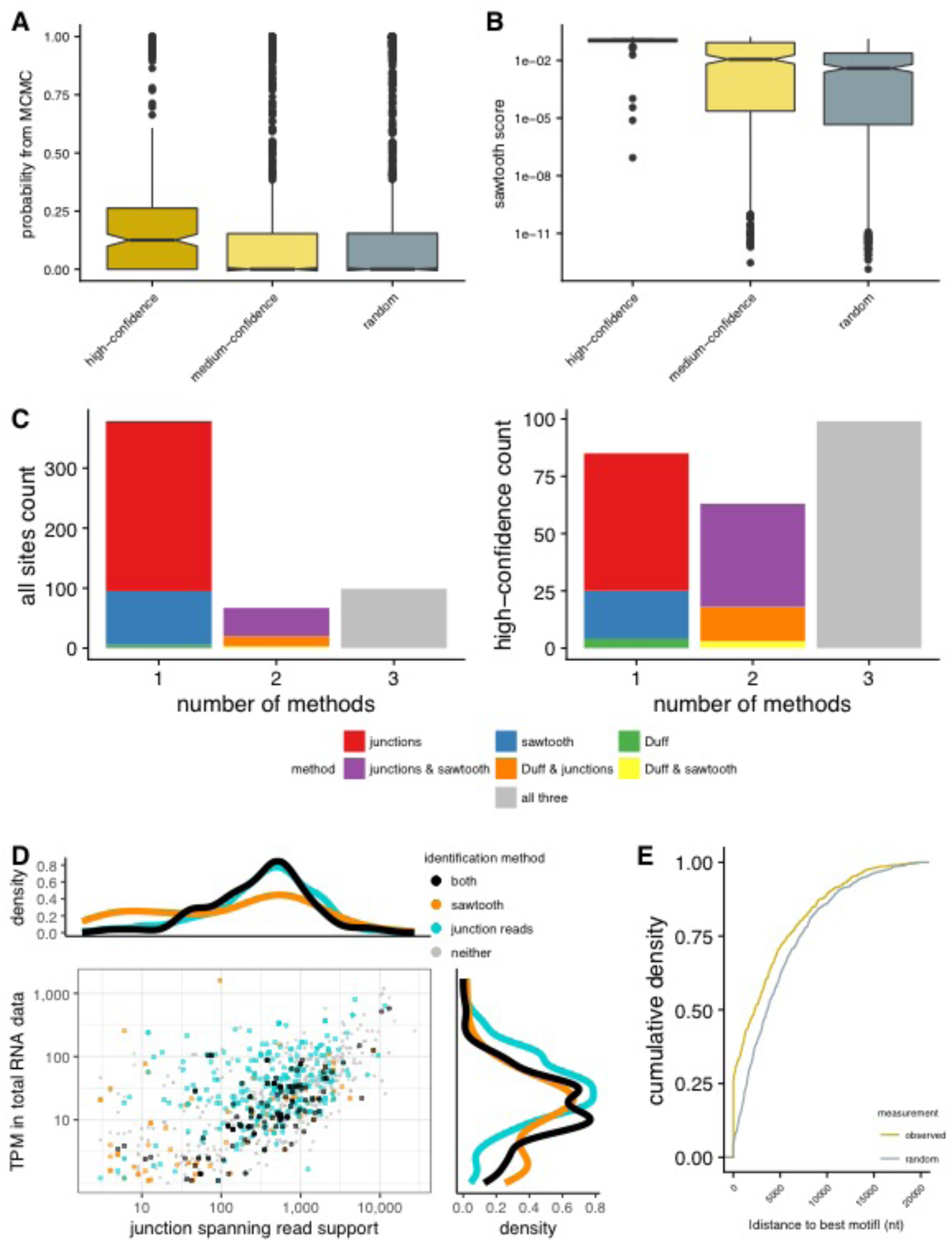
Identifying sites of recursive splicing. **(A)** The probability derived from the sawtooth MCMC model of a site being a recursive site for the final set of recursive sites (*light orange*), all sites with minimal support from any method (*dark orange*), and random sites placed down in the same introns (*grey*). **(B)** The sawtooth score (see Methods) for the final set of recursive sites (*light orange*), all sites with minimal support from any method (*dark orange*), and random sites place down in the same introns (*grey*). **(C)** Number of recursive sites (*left*) and high-confidence sites (*right*) identified by one of multiple identification pipelines, with the majority of recursive sites identified by both junction reads and sawtooth scores, as well as present in the Duff *et al*. dataset. **(D)** The gene expression levels of genes with recursive introns (TPM, *y-axis*) relative to the junction spanning read support for each recursive intron (read count, *x-axis*), showing the varying power to identify recursive sites with the sawtooth recursive method (*orange*), junction-spanning reads alone (*blue*), or both methods (*black*). **(E)** The cumulative distribution of distances between the recursive site identified with the sawtooth recursive method and the best matching recursive motif (orange) and random sites placed down in the same introns (grey) are significantly different.

**Figure 2 – figure supplement 1.**
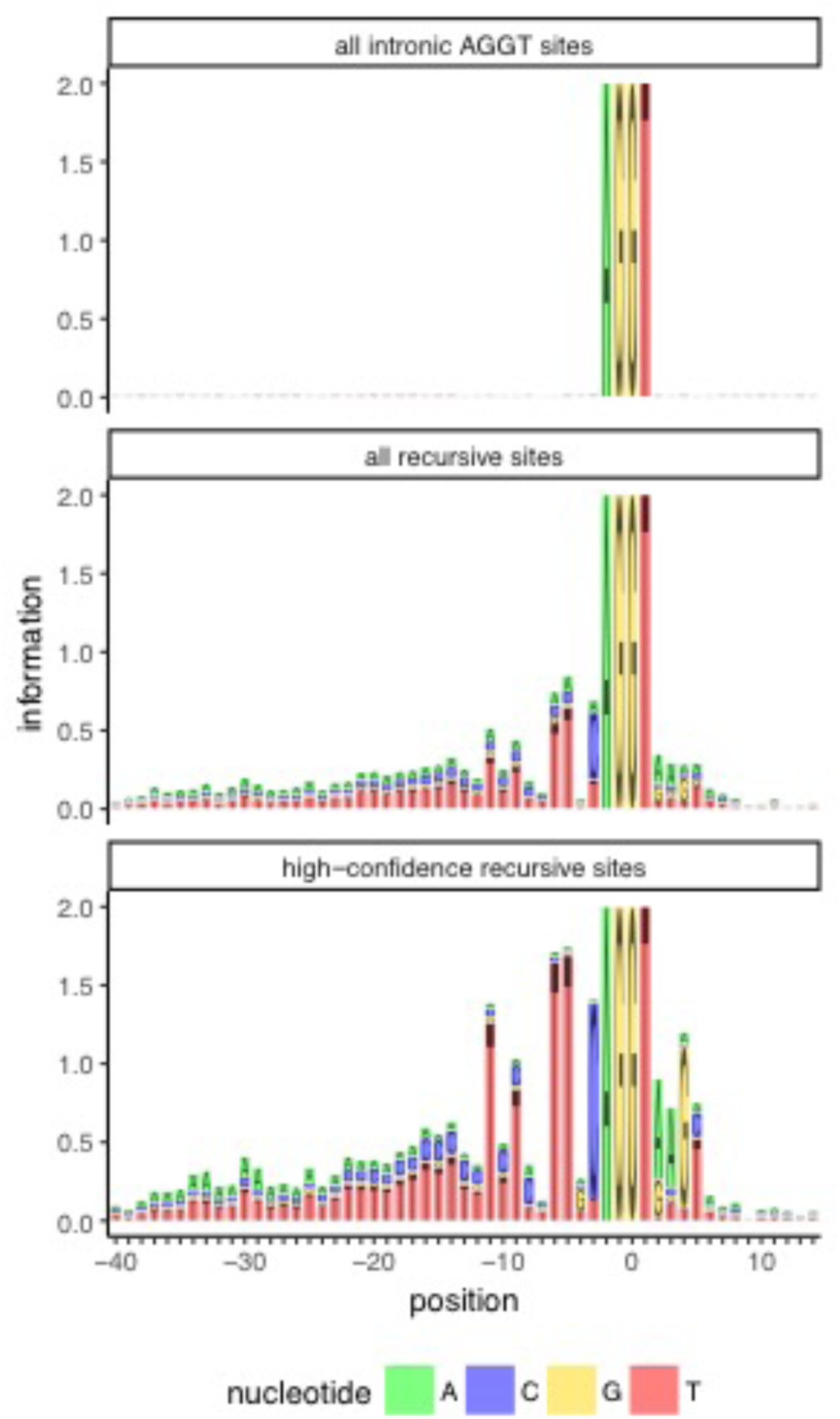
Properties of recursively spliced introns. Sequence logo for all intronic AG|GT sites (*top*), medium-confidence recursive sites (*middle*) and high-confidence recursive sites (*bottom*).

**Figure 4 – figure supplement 1.**
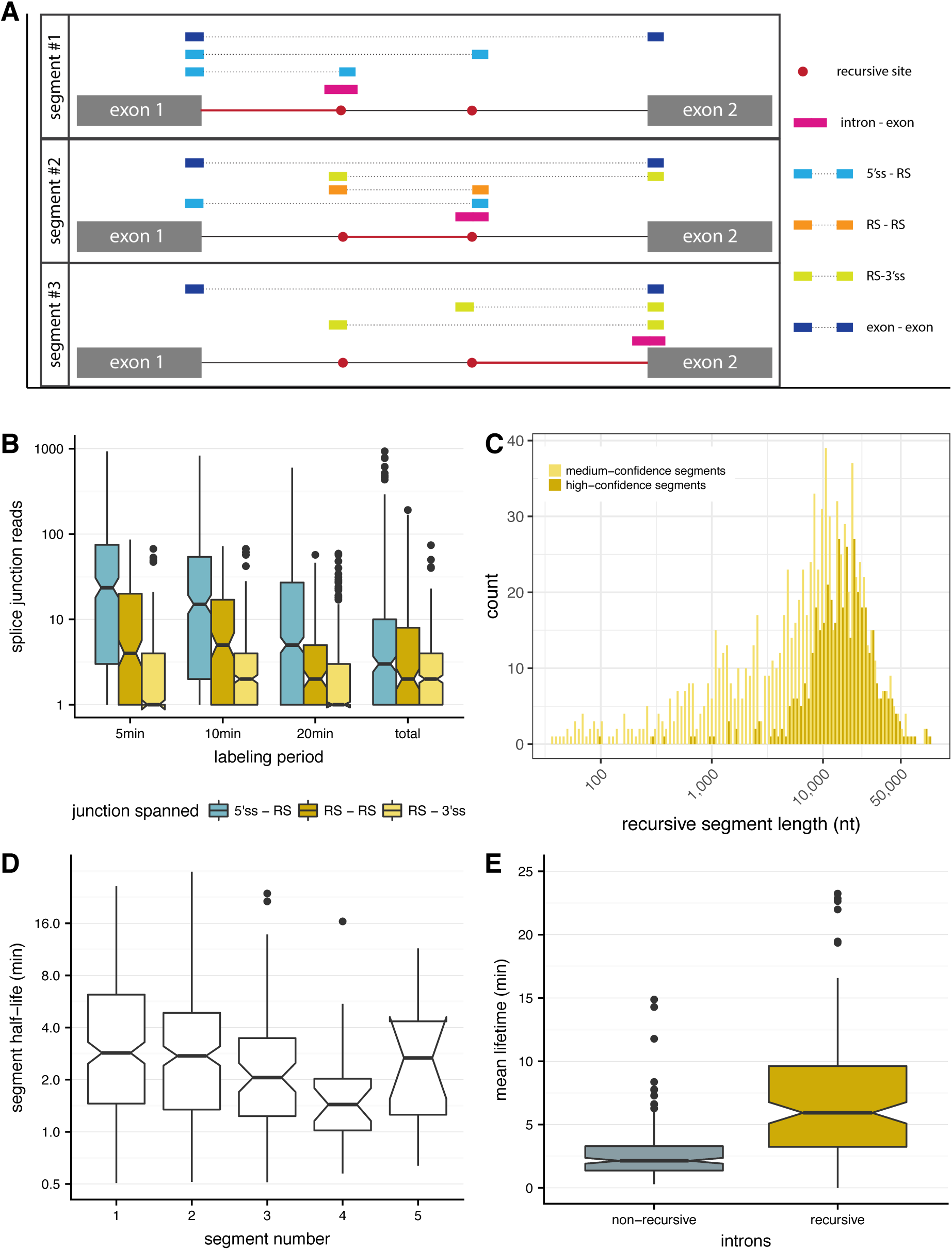
Rates of recursive splicing. **(A)** Junction reads used to estimate splicing half-lives for recursive segments (*red lines*), centered on 3’ recursive sites (*red dots*) for each segment. Incomplete splicing is estimated from intron-exon junction reads (*pink bars*). Completed splicing is estimated from a sum across split-junction reads between the 5’ splice site and recursive site (*light blue bars*), two recursive sites (*orange bars*), a recursive site and the 3’ splice site (*yellow bars*), and the 5’ splice site and 3’ splice site (exon-exon read, *dark blue bars*). Each segment’s splicing is informed by different types of junction reads dependent on the position in the intron, as drawn for an intron with three recursive segments. **(B)** The number of splice junction reads (y-axis) spanning a 5’ splice site and recursive site (*blue*), two recursive sites (*gold*), and a recursive site and 3’ splice site (*yellow*) across the labeling periods (*x-axis*). **(C)** Distribution of lengths of recursive segments (nucleotides, *x-axis*) for medium-confidence recursive segments (*yellow*) and high-confidence recursive segments (*gold*). **(D)** Splicing half-lives (*y-axis*) for recursive segments with varying positions across the intron (*x-axis*), where on average, all segments in an intron tend to be spliced out at similar rates. **(E)** The distribution of mean life-times (y-axis) for recursively spliced introns (estimated by the maximum of exponentials from constituent recursive segment splicing rates, *gold*) relative to non-recursive introns chosen to match the length of the recursive introns (*grey*).

**Figure 4 – figure supplement 2.**
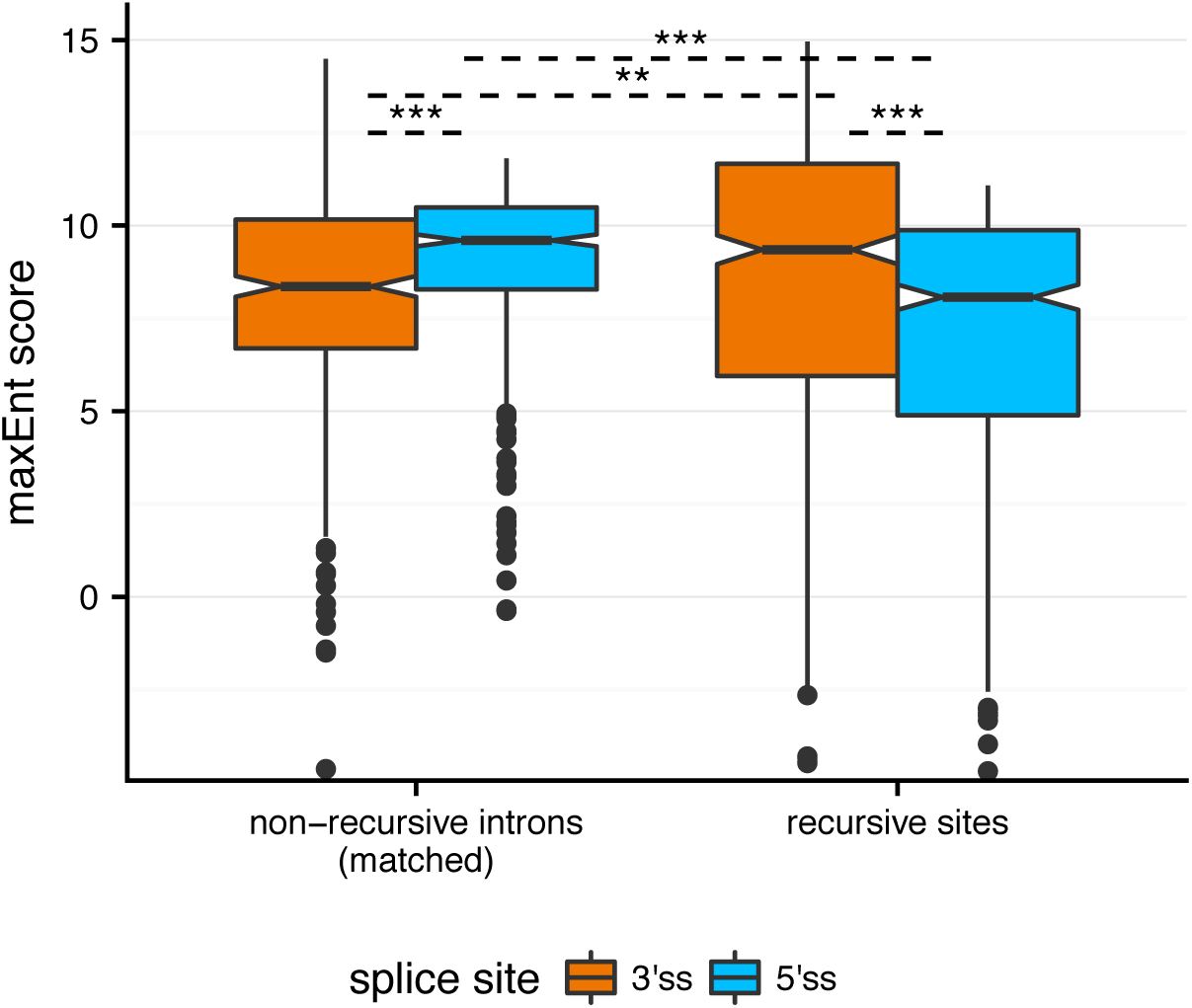
Recursive splice site motif strength. Distribution of splice site strengths (maxEnt score, *y-axis*) across both 3’ splice sites (*orange*) and 5’ splice sites (*blue*) for recursive sites (*right*) and non-recursive introns matched for intron length (*left*).Significance is indicated such that **: P < 0.01 and ***: P < 0.001, with a Mann-Whitney U test.

### Supplementary File 1: Summary statistics and information for recursive sites

*Column 1 – intron:* Coordinates of intron containing recursive site, with chr:start-end:strand.

*Column 2 – gene:* FlyBase gene symbol for parent gene.

*Column 3 – TPM:* Gene expression values calculated using kallisto (TPMs).

*Column 4 – completed_splicing_junction_reads:* Number of junction reads supporting completed splicing across the entire intron.

*Column 5 – recursive_site:* Coordinate for the recursive site.

*Column 6 – method:* Method used for identification of the recursive site, where “junction” indicates site identified by either *RachetJunction* or *RachetPair*, “sawtooth” indicates site identified by *RachetScan*, and “both” indicates site identified by both methods.

*Column 7 – in_duff:* Flag indicating the recursive site was identified in the Duff *et al*. study.

*Column 8 – high_confidence:* Flag indicating the recursive site was identified as a high-confidence site (1) or a medium-confidence site (0).

*Column 9 – junction_reads:* Comma-separated list of the number of junction reads (5’-RS) supporting the recursive site in each timepoint (combined across replicates) [5m, 10m, 20m, total].

*Column 10 – spanning_read_pairs:* Comma-separated list of number of spanning read-pairs supporting the recursive site in each timepoint (combined across replicates) [5m, 10m, 20m, total].

*Column 11 – sawtooth_score:* Sawtooth score for the recursive site, as defined in the Methods.

*Column 12 – mcmc_probability:* Probability of this site being a recursive site, as derived from the MCMC sampling procedure integral to the *RachetScan* method.

*Column 13 – recursive_index:* Recursive index for the recursive site, as defined in the Methods.

*Column 14 – motif:* Sequence found around the recursive site.

*Column 15 – motif_score*. Motif score for the recursive site, as defined in the Methods.

*Column 16 – downstream_reads:* Number of splice junction reads originating from the 3’ end of the exon.

*Column 17 – intron_body_reads:* Number of reads in the body of the intron.

**Supplementary File 2: Gene ontology enrichment for all recursive sites and high-confidence recursive sites**.

**Supplementary File 3: Summary statistics and information about rates of recursive spliced segments**.

*Column 1 – intron:* Coordinates of intron containing recursive site, with chr:start-end:strand.

*Column 2 – gene:* FlyBase gene symbol for parent gene.

*Column 3 – recursive_site:* Coordinate for the recursive site (or 3’ splice site in the case of the final segment of the intron).

*Column 4 – segment_type:* Indicator whether the segment is spliced to a recursive site. (“segment”) or to the 3’ splice site (“threess”, in the case of the final segment of the intron).

*Column 5 – segment_len:* Length of the segment (nucleotides).

*Column 6 – segment_num:* Position of the segment relative to other segments in the intron.

*Column 7 – three_length:* Length of the region from the recursive site (or 3’ splice site) to the polyA site of the transcript (nucleotides).

*Columns 8-10 – ie_count_[timepoint]:* count of the intron-exon junction reads for each of the labeling periods (summed across three replicates per labeling period) – for recursive sites, this overlaps the recursive site, while for the final segment this overlaps the 3’ splice site.

*Columns 11-13 – ee_count_[timepoint]:* count of the exon-exon junction reads for each of the labeling periods (summed across three replicates per labeling periods) – this includes junctions deriving from recursive intermediates, as outlined in the Methods.

*Column 14 – halflife:* Half-life of the recursive segment computed using the junction dynamics approach described in Pai *et al*. 2017 (min).

*Column 15 – txn_to_three:* Time to transcribe the remainder of the intron from the recursive site to the 3’ splice site (min).

## References

1. Hatton AR, Subramaniam V, Lopez AJ. Generation of Alternative Ultrabithorax Isoforms and Stepwise Removal of a Large Intron by Resplicing at Exon–Exon Junctions. Molecular Cell. 1998;2: 787–796. doi:10.1016/S1097-2765(00)80293-2

2. Conklin JF, Goldman A, Lopez AJ. Stabilization and analysis of intron lariats in vivo. Methods. 2005;37: 368–375. doi:10.1016/j.ymeth.2005.08.002

3. Burnette JM, Miyamoto-Sato E, Schaub MA, Conklin J, Lopez AJ. Subdivision of Large Introns in Drosophila by Recursive Splicing at Nonexonic Elements. Genetics. Genetics; 2005;170: 661–674. doi:10.1534/genetics.104.039701

4. Duff MO, Olson S, Wei X, Garrett SC, Osman A, Bolisetty M, et al. Genome-wide identification of zero nucleotide recursive splicing in Drosophila. Nature. Nature Research; 2015;521: 376–379. doi:10.1038/nature14475

5. Sibley CR, Emmett W, Blazquez L, Faro A, Haberman N, Briese M, et al. Recursive splicing in long vertebrate genes. Nature. Nature Publishing Group; 2015;521: 371–375. doi:10.1038/nature14466

6. Singh J, Padgett RA. Rates of in situ transcription and splicing in large human genes. Nature Structural & Molecular Biology. Nature Publishing Group; 2009;16: 1128–1133. doi:10.1038/nsmb.1666

7. Schmidt U, Basyuk E, Robert M-C, Yoshida M, Villemin J-P, Auboeuf D, et al. Real-time imaging of cotranscriptional splicing reveals a kinetic model that reduces noise: implications for alternative splicing regulation. J Cell Biol. Rockefeller University Press; 2011;193: 819–829. doi:10.1083/jcb.201009012

8. Martin RM, Rino J, Carvalho C, Kirchhausen T, Carmo-Fonseca M. Live-Cell Visualization of Pre-mRNA Splicing with Single-Molecule Sensitivity. Cell Reports. 2013;4: 1144–1155. doi:10.1016/j.celrep.2013.08.013

9. Pai AA, Henriques T, McCue K, Burkholder A, Adelman K, Burge CB. The kinetics of pre-mRNA splicing in theDrosophilagenome and the influence of gene architecture. eLife Sciences. eLife Sciences Publications Limited; 2017;6: 1123. doi:10.7554/eLife.32537

10. Guo Y, Mahony S, Gifford DK. High resolution genome wide binding event finding and motif discovery reveals transcription factor spatial binding constraints. Aerts S, editor. PLoS Comp Biol. Public Library of Science; 2012;8: e1002638. doi:10.1371/journal.pcbi.1002638

11. Brugiolo M, Herzel L, Neugebauer KM. Counting of co-transcriptional splicing. F1000Prime Rep. 2013.

12. Tadros W, Lipshitz HD. The maternal-to-zygotic transition: a play in two acts. Development. The Company of Biologists Ltd; 2009;136: 3033–3042. doi:10.1242/dev.033183

13. Artieri CG, Fraser HB. Transcript Length Mediates Developmental Timing of Gene Expression Across Drosophila. Molecular Biology and Evolution. Oxford University Press; 2014;31: 2879–2889. doi:10.1093/molbev/msu226

14. Bray NL, Pimentel H, Melsted P, Pachter L. Near-optimal probabilistic RNA-seq quantification. Nature Biotechnology. Nature Publishing Group; 2016. doi:10.1038/nbt.3519

15. St Pierre SE, Ponting L, Stefancsik R, McQuilton P, FlyBase Consortium. FlyBase 102— advanced approaches to interrogating FlyBase. Nucleic Acids Research. Oxford University Press; 2014;42: D780–D788. doi:10.1093/nar/gkt1092

16. Siepel A, Bejerano G, Pedersen JS, Hinrichs AS, Hou M, Rosenbloom K, et al. Evolutionarily conserved elements in vertebrate, insect, worm, and yeast genomes. Genome Research. 2005;15: 1034–1050. doi:10.1101/gr.3715005

17. Kim D, Langmead B, Salzberg SL. HISAT: a fast spliced aligner with low memory requirements. Nat Meth. 2015;12: 357–360. doi:10.1038/nmeth.3317

18. Heger A, Jacobs K. pysam: htslib interface for python [Internet]. [cited 26 Feb 2018]. Available: https://github.com/pysam-developers/pysam

19. Smit A, Hubley R, Green P. RepeatMasker Open-4.0 [Internet]. [cited 27 Feb 2018]. Available: http://www.repeatmasker.org

20. Jones E, Oliphant E, Peterson P. Scipy: Open Source Scientific Tools for Python [Internet]. [cited 27 Feb 2018]. Available: http://www.scipy.org

21. Zare-Mirakabad F, Ahrabian H, Sadeghi M, Nowzari-Dalini A, Goliaei B. New scoring schema for finding motifs in DNA Sequences. BMC Bioinformatics. 2009;10: 93. doi:10.1186/1471-2105-10-93

22. Li H, Handsaker B, Wysoker A, Fennell T, Ruan J, Homer N, et al. The Sequence Alignment/Map format and SAMtools. Bioinformatics. 2009;25: 2078–2079. doi:10.1093/bioinformatics/btp352

23. Dobin A, Davis CA, Schlesinger F, Drenkow J, Zaleski C, Jha S, et al. STAR: ultrafast universal RNA-seq aligner. Bioinformatics. Oxford University Press; 2013;29: 15–21. doi:10.1093/bioinformatics/bts635

24. Yeo G, Burge CB. Maximum Entropy Modeling of Short Sequence Motifs with Applications to RNA Splicing Signals. http://www.liebertpubcom/cmb. Mary Ann Liebert, Inc; 2004. doi:10.1089/1066527041410418

25. Gene Ontology Consortium. Gene Ontology Consortium: going forward. Nucleic Acids Research. 2015;43: D1049–56. doi:10.1093/nar/gku1179

